# An experimentally evolved variant of RsmA confirms its central role in the control of *Pseudomonas aeruginosa* social motility

**DOI:** 10.1101/2020.07.15.203992

**Authors:** Sophie Robitaille, Yossef López de los Santos, Marie-Christine Groleau, Fabrice Jean-Pierre, Nicolas Doucet, Jonathan Perreault, Eric Déziel

**Author notes:** Department of Microbiology and Immunology, Geisel School of Medicine at Dartmouth, Hanover, New Hampshire, USA.

## Abstract

Bacteria can colonize a variety of different environments by modulating their gene regulation using two-component systems. The versatile opportunistic pathogen *Pseudomonas aeruginosa* has been studied for its capacity to adapt to a broad range of environmental conditions. The Gac/Rsm pathway is composed of the sensor kinase GacS, that detects environmental cues, and the response regulator GacA, that modulates the expression of a specific genes. This system, through the sRNA repressors RsmY and RsmZ, negatively controls the activity of the protein RsmA, which is centrally involved in the transition from chronic to acute infections by post-transcriptionally regulating several virulence functions. RsmA positively regulates swarming motility, a social surface behaviour. Through a poorly defined mechanism, RsmA is also indirectly regulated by HptB, and a Δ*hptB* mutant exhibits a severe swarming defect. Since a Δ*hptB* mutant retains all the known functions required for that type of motility, we used an experimental evolution approach to identify elements responsible for its swarming defect. After a few passages under swarming conditions, the defect of the Δ*hptB* mutant was rescued by the emergence of spontaneous single nucleotide substitutions in the *gacA* and *rsmA* genes. Since GacA indirectly represses RsmA activity, it was coherent that an inactivating mutation in *gacA* would compensate the Δ*hptB* swarming defect. However, the effect of the mutation in *rsmA* was unexpected since RsmA promotes swarming; indeed, using expression reporters, we found that the mutation that does not abolish its activity. Instead, using electrophoretic mobility shift assays and molecular simulations, we show that this variant of RsmA is actually less amenable to titration by its cognate repressor RsmY, supporting the other phenotypes observed for this mutant. These results confirm the central role of RsmA as a regulator of swarming motility in *P. aeruginosa* and identify residues crucial for RsmA function in social motility.

**Author summary:** Bacteria need to readily adapt to their environment. Two-component systems (TCS) allow such adaption by triggering bacterial regulation changes through the detection of environmental cues. The opportunistic pathogen *Pseudomonas aeruginosa* possesses more than 60 TCS in its genome. The Gac/Rsm is a TCS extensively studied for its implication in virulence regulation. This system regulates the transition between chronic and acute bacterial infection behaviours. To acquire a better understanding of this regulation, we performed a directed experimental evolution on a swarming-deficient mutant in a poorly understood regulatory component of the Gac/Rsm pathway. We observed single nucleotide substitutions that allowed restoration of a swarming phenotype similar to the wild-type behaviour. More specifically, mutations were found in the *gacA* and *rsmA* genes. Interestingly, the observed mutation in *rsmA* does not result in loss of function of the protein but rather alters its susceptibility to repression by its cognate interfering sRNA. Since modification in the RNA sequence of RsmA results in the rescue of swarming motility, we confirm the central role of this posttranscriptional repressor in this social lifestyle.

## Introduction

Bacteria can adapt to diverse environments using various mechanisms. They use two-component systems (TCS) to rapidly modulate the expression of specific subsets of genes (1). TCS convert external stimuli into an internal response that promotes adaptation to environmental cues. Some bacteria exploit these systems for virulence regulation (2). Typically, TCS consists of a histidine sensor kinase that responds to an external signal to trigger the autophosphorylation of an intracellular histidine residue. Then, the phosphoryl group of the sensor kinase is transferred to an aspartate residue located in the receiver domain of a cognate response regulator, which then modulates the expression of a specific set of target genes (3). In some cases, phosphorylation of the receiver domain can occur through a His-containing phosphotransfer (Hpt) protein that acts as an intermediate between the membrane sensor and the response regulator (4).

*Pseudomonas aeruginosa* is an opportunistic pathogen responsible for several nosocomial infections and also a major cause of morbidity and mortality among individuals with cystic fibrosis (5, 6). The genome of prototypical *P. aeruginosa* strain PAO1 contains 63 histidine kinases, 64 response regulators, and three Hpt proteins (7). The Gac/Rsm pathway regulates, among others, virulence-associated genes, and biofilm formation (8). This pathway regulates the transition between chronic (associated with the sessile lifestyle) and acute (associated with the motile lifestyle) infections (9, 10). The Gac TCS is composed of the histidine sensor kinase GacS (11) and its cognate response regulator GacA. When GacS is phosphorylated, it transfers its phosphoryl group to GacA (12), promoting the transcription of the small RNAs (sRNA) RsmY and RsmZ (10). The expression levels of these sRNAs are influenced by bacterial culture conditions, either in broth or on a surface, highlighting the importance of this pathway in the regulation between these two modes of growth (13). HptB modulates the expression of RsmZ through an unknown surface-specific membrane sensor other than GacS, resulting in the phosphorylation of GacA and/or an uncharacterized regulatory factor (13). The sRNAs RsmY and RsmZ are repressors of the protein RsmA and act by reversibly titrating its activity (14–16). RsmA is a post-transcriptional regulator that binds to a specific trinucleotide GGA motif usually present in the 5’UTR of target mRNAs, thus preventing their translation (8, 17). RsmA inhibits genes associated with biofilm development and favors genes linked to acute infections (10). Also, RsmA is necessary for colonization in a murine infection model (18).

Swarming motility is a social surface motility behaviour utilized by several bacteria to promote the colonization of environments by coordinating the movement of a bacterial population on a semi-solid surface (19). Swarming cells require a functional rotating flagellum which gives them the necessary propelling power. They also need to produce surface-active agents reducing the surface tension between the substrate and the cell, thus facilitating the movement on the surface (19). Loss of RsmA or HptB functions leads to defects in swarming motility (13, 20, 21), highlighting the importance of this regulatory cascade in swarming regulation.

To date, no specific gene directly regulated by RsmA has been identified to explain the control of swarming motility. We have previously determined that the defect in swarming motility of a Δ*hptB* mutant is not due to either a lack of functional flagella or a deficit in surfactant production (13). Therefore, additional and still unknown elements are necessary for the swarming motility of *P. aeruginosa.* Since swarming motility is believed to be a beneficial phenotype, defective mutants should derive a selective benefit to regain social motility behaviour. Indeed, Boyle *et al.* (22) rescued the swarming motility of a *cbrA* mutant through directed experimental evolution. To test our hypothesis, we recently performed a similar experiment using directed swarming evolution on a Δ*hptB* mutant of strain PA14 to identify unknown elements necessary for swarming motility (23). After four transfers corresponding to cell passages from the tips of swarming tendrils on new swarming plates, the swarming motility of Δ*hptB* was restored (23). Among evolved gain-of-function clones, two distinct phenotypes emerged: (i) pyocyanin-overproducers with a partially restored swarming phenotype resulting from mutations in *lasR* (described in (23)); and (ii) clones expressing a completely restored swarming behaviour comparable to the parental strain PA14. The latter clones (named C2 and C4) are described and characterized here (Fig. 1 B-C); we found single nucleotide substitutions in the *rsmA* and *gacA* genes, further strengthening the importance of the Gac/Rsm pathway in the regulation of surface social behaviour in *P. aeruginosa.* In particular, this experimental evolution experiment revealed residues crucial for RsmA interaction with RNA.

**Figure 1.**
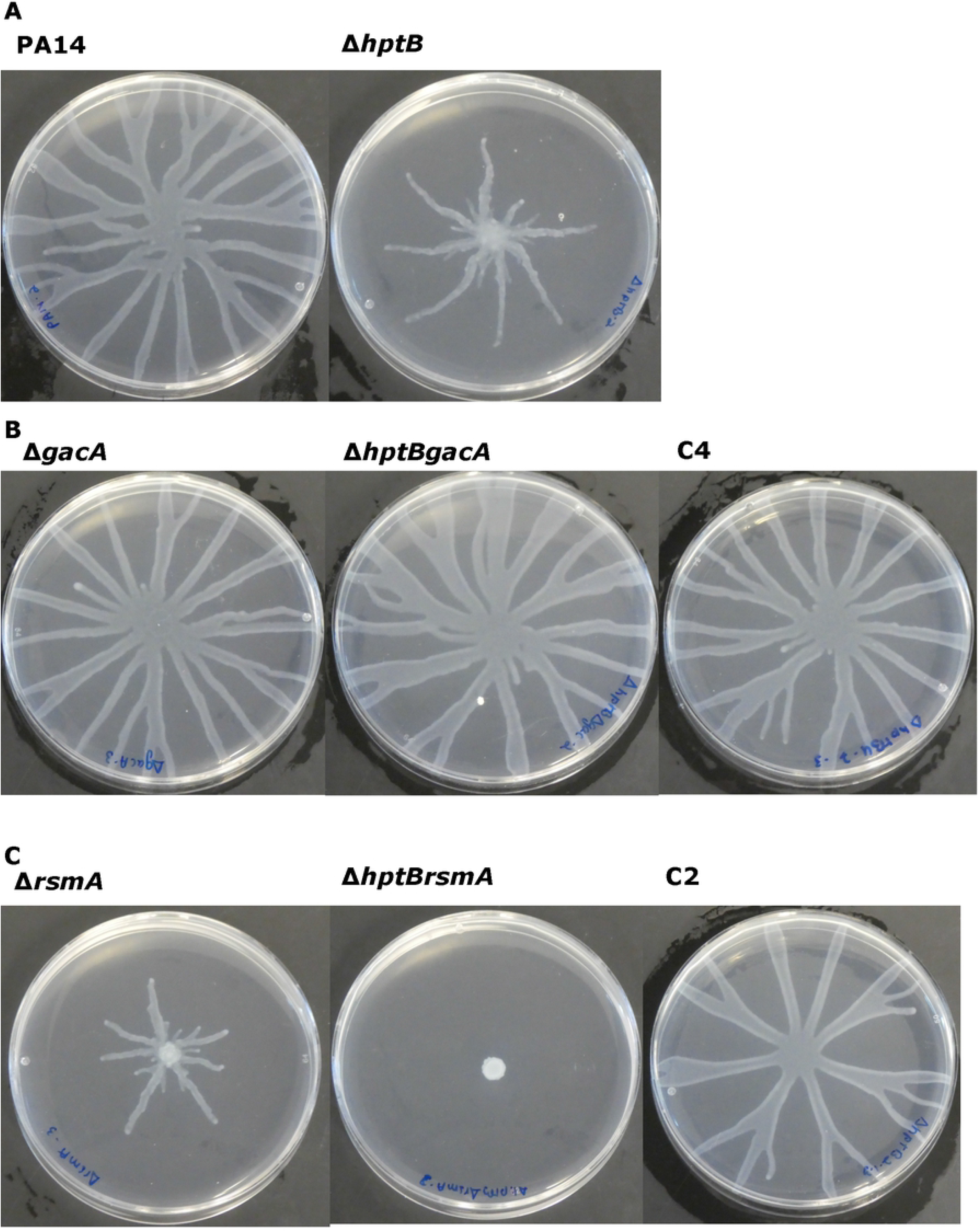
Swarming phenotype of the C2 and C4 clones is similar to wild-type *Pseudomonas aeruginosa* PA14. Swarming motility of (A) PA14 and Δ*hptB* (B) *△gacA, ΔhptBgacA,* and clone C4, (C) Δ*rsmA, ΔhptBrsmA,* clone C2 after 20h of incubation at 34°C.

## Results

### Evolution of the Δ*hptB* mutant restores swarming phenotype by selecting mutations in the Gac/Rsm regulatory pathway

We previously reported a directed swarming evolution experiment on the Δ*hptB* mutant of *P. aeruginosa* PA14 to identify elements capable of restoring a swarming phenotype similar to wild-type (23). Two different groups of clones had partially or completely restored swarming abilities. Thus, the evolved Δ*hptB* clones acquired compensatory mutations. We previously reported that mutations in *lasR* characterized the partially recovered mutants (23). Here, we focus on mutations involved in the complete recovery of swarming motility of the Δ*hptB* mutant. Whole-genome sequencing was performed on clones C2 and C4, which completely regained their swarming phenotype (**Fig. 1**) (23). Clone C4 has a single non-synonymous point mutation in the *gacA* gene (PA14_30650), resulting in a substitution at residue 90 (c.268C>T; proline to serine (p.P90S)) when compared to the parental Δ*hptB* protein sequence. Clone C2 carries a mutation (c.91C>A) in the *rsmA* gene (PA14_52570), resulting in an arginine to serine substitution at position 31 of the protein sequence (p.R31S) when compared to the parental Δ*hptB* genome sequence. No additional mutations were identified in these two clones.

Given the emergence of non-synonymous mutations in the *gacA* and *rsmA* genes (clones C4 and C2, respectively) after repeated passages of the Δ*hptB* isogenic strain to select for a recovered swarming phenotype, we verified whether the acquired mutations were responsible for the rescue. We first looked at the activity of GacA in the C4 clone. As shown in **figure 1B**, inactivation of *gacA* in a Δ*hptB* strain restores the swarming phenotype. Accordingly, swarming of clone C4 is similar to that of Δ*gacA* and Δ*hptBgacA* mutants. Furthermore, we verified whether the activity of GacA is affected in clone C4 by looking at the transcription of its direct targets, *rsmY* and *rsmZ* (**Fig. 2 and Fig. S1**). In agreement with a loss of GacA function, the transcription of both sRNAs is significantly lower in the evolved clone when compared to its parental Δ*hptB* background. Clone C4 shows no significant difference in expression of *rsmY* and *rsmZ* with the Δ*gacA* and Δ*hptBgacA* mutants. Thus, these results demonstrate that clone C4 acquired an inactivating mutation in the GacA regulator, allowing for the rescue of the swarming defect in the Δ*hptB* parental strain.

**Figure 2.**
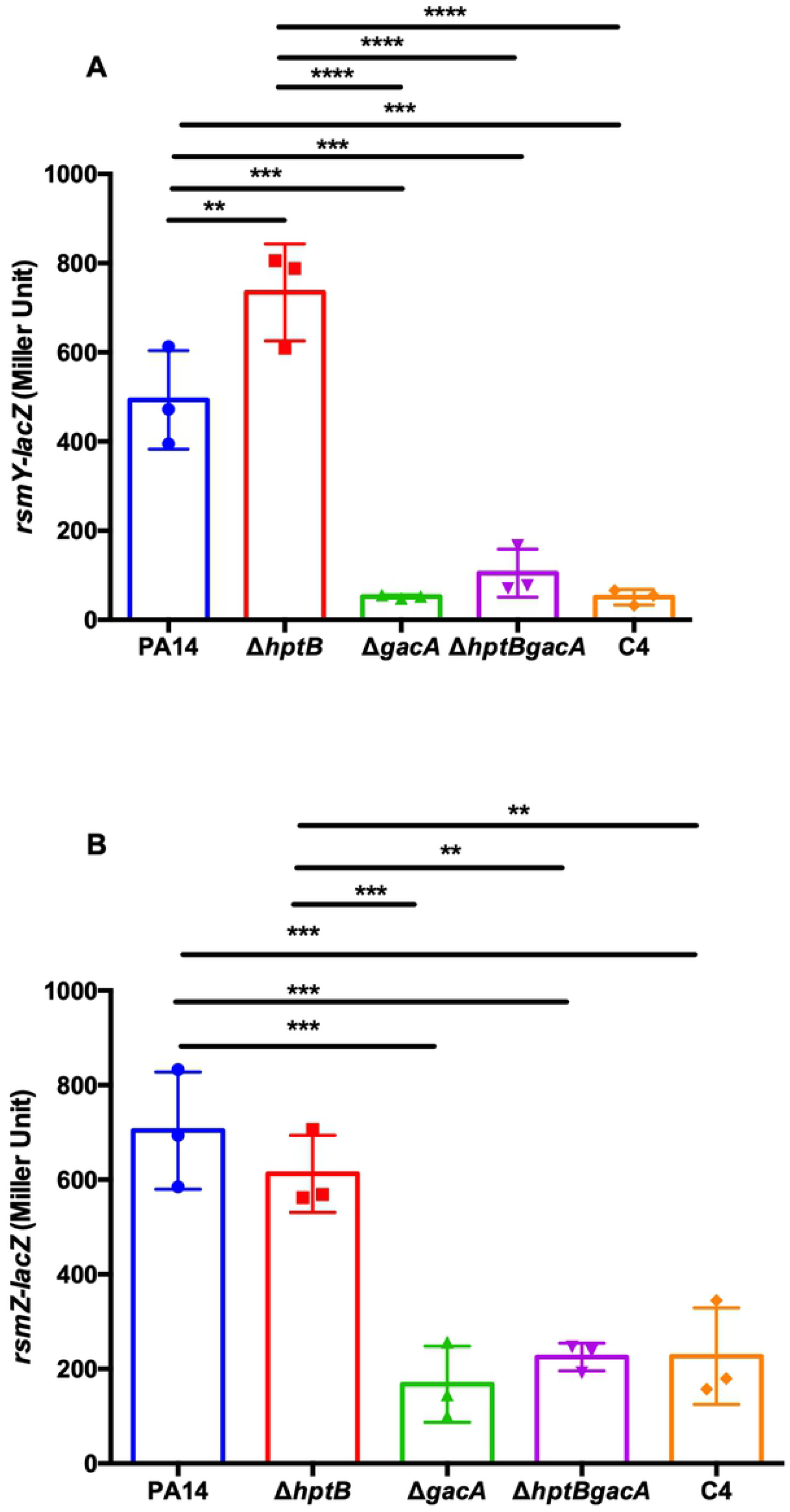
Analysis of *rsmY* and *rsmZ* expression in the C4 clone mutant. (A) β-galactosidase activity of a *rsmY-lacZ* reporter (B) and *rsmZ-lacZ* reporter in bacteria grown in TSB at 37°C after 5h of incubation. One-way ANOVA was performed comparing each strain to each other (*≤0.05, **≤0.01, ***≤0.001, ****≤0.0001). See **Fig. S1** for the growth of each strain.

We then looked at the effect of the *rsmA* mutation in the C2 clone. Loss of *hptB* or *rsmA* usually results in an important defect in swarming motility (**Fig. 1A-C**) (13,21,24), while a combination of both *rsmA* and *hptB* mutations results in a complete loss of coordinated social movement (**Fig. 1C**). However, swarming was rescued in clone C2 (**Fig. 1C**). This was surprising given that the over-expression of RsmA in the Δ*hptB* background also results in a rescue of the swarming phenotype (**Fig. S2**). Thus, we considered two possible explanations for these results: [1] the single nucleotide substitution in *rsmA* does not influence the function of RsmA, and another undetected mutation is responsible for the swarming recovery, or [2] the arginine-to-serine substitution in RsmA in the evolved clone C2 somehow increases the activity of this regulator. To verify the first possibility, which we thought very unlikely considering the high coverage of our genome sequencing, we looked at the complementation of a Δ*rsmA* markerless mutant with a plasmid carrying the RsmA^R31S^ substitution from clone C2 by measuring the transcription of *rsmY.* The transcription of *rsmY* and *rsmZ* are downregulated in a Δ*rsmA* mutant background (15, 24). We observed that a plasmid-borne *rsmA*^R31S^ can restore expression of *rsmY* in Δ*rsmA,* although incompletely (**Fig. 3 and Fig. S3**). Thus, RsmA^R31S^ is functional, but with a somewhat altered activity, indicating that the mutation in clone C2 does not result in abrogation of the activity of the protein. We then looked at the translation of *hcnA* mRNA transcripts, known to be directly repressed by RsmA (25, 26). Interestingly, we observed that the translation of *hcnA* is lower in the C2 clone compared to the Δ*hptBrsmA* mutant, but similar to the wild-type strain (**Fig. 4A and Fig.S4A**). Thus, to further understand the impact of the observed non-synonymous mutation in the *rsmA* gene of clone C2, we looked at the transcription of *rsmY* and *rsmZ* (**Fig. 4 B-C and Fig.S4B-C**). We observed that the C2 clone exhibits a higher expression of both sRNAs when compared to the Δ*rsmA* and Δ*hptBrsmA* mutants, while it is lower than in the Δ*hptB* mutant. These results confirm that RsmA is still functional in clone C2. In fact, the RsmA regulatory activity of the evolved clone is similar to that of the wild-type PA14 strain (**Fig. 4 A-C**). Taken together, these data indicate that RsmA^R31S^ is active in the C2 clone, but its function is altered when compared to wild-type RsmA. Importantly, RsmA^R31S^ is able to rescue the swarming defect imposed by the loss of HptB.

**Figure 3.**
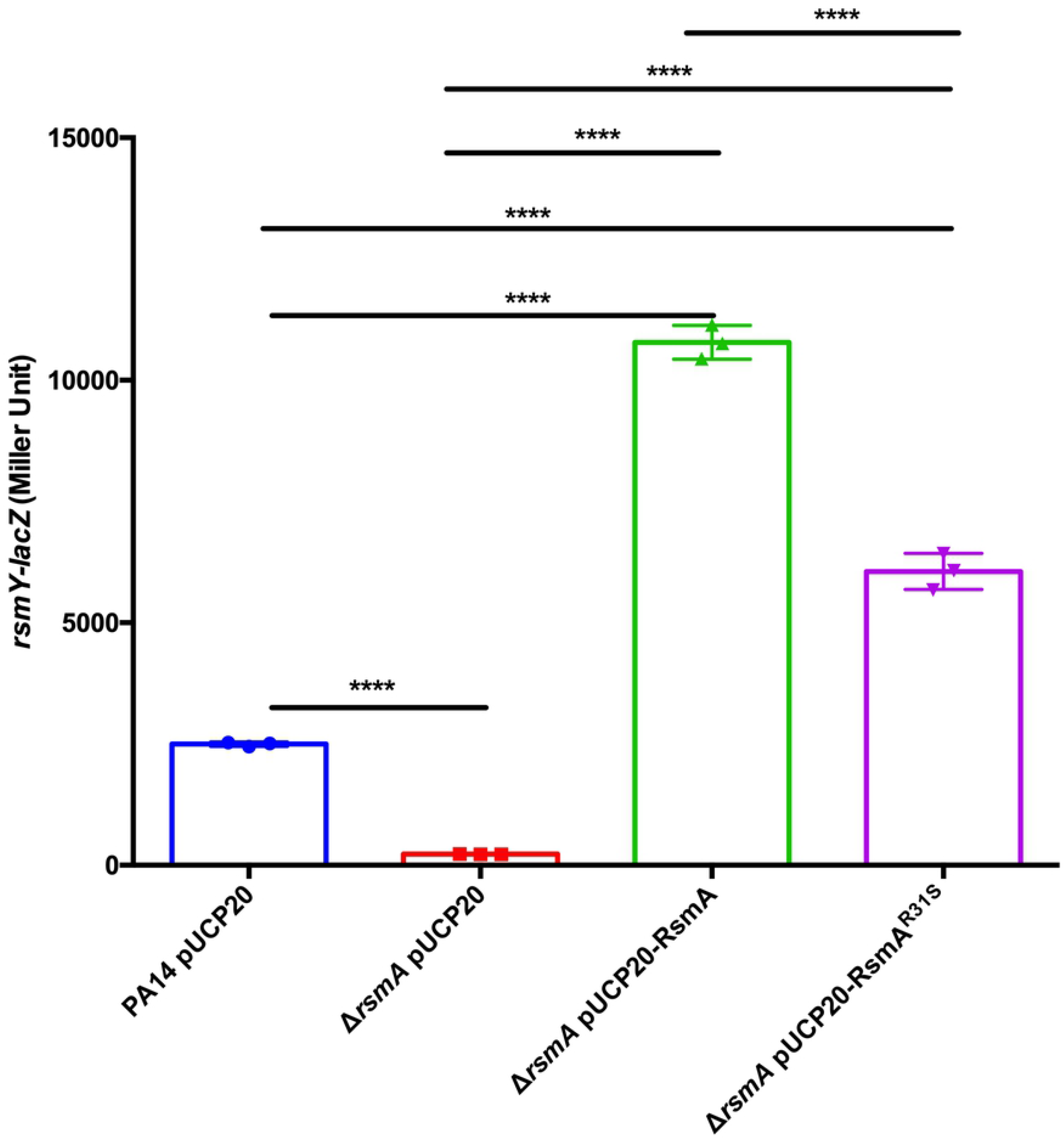
Complementation of mutant *rsmA* with pUCP20 as a control, pSR15 (pUCP20-*rsmA*) and pSR16 (pUCP20-*rsmA*^R31S^) for the *rsmY-*lacZ expression at 6h in TSB at 37°C. One-way ANOVA was performed to compare each strain to each other. (*≤0.05, **≤0.01, ***≤0.001, ****≤0.0001). See **Fig. S3** for the growth of each strain.

**Figure 4.**
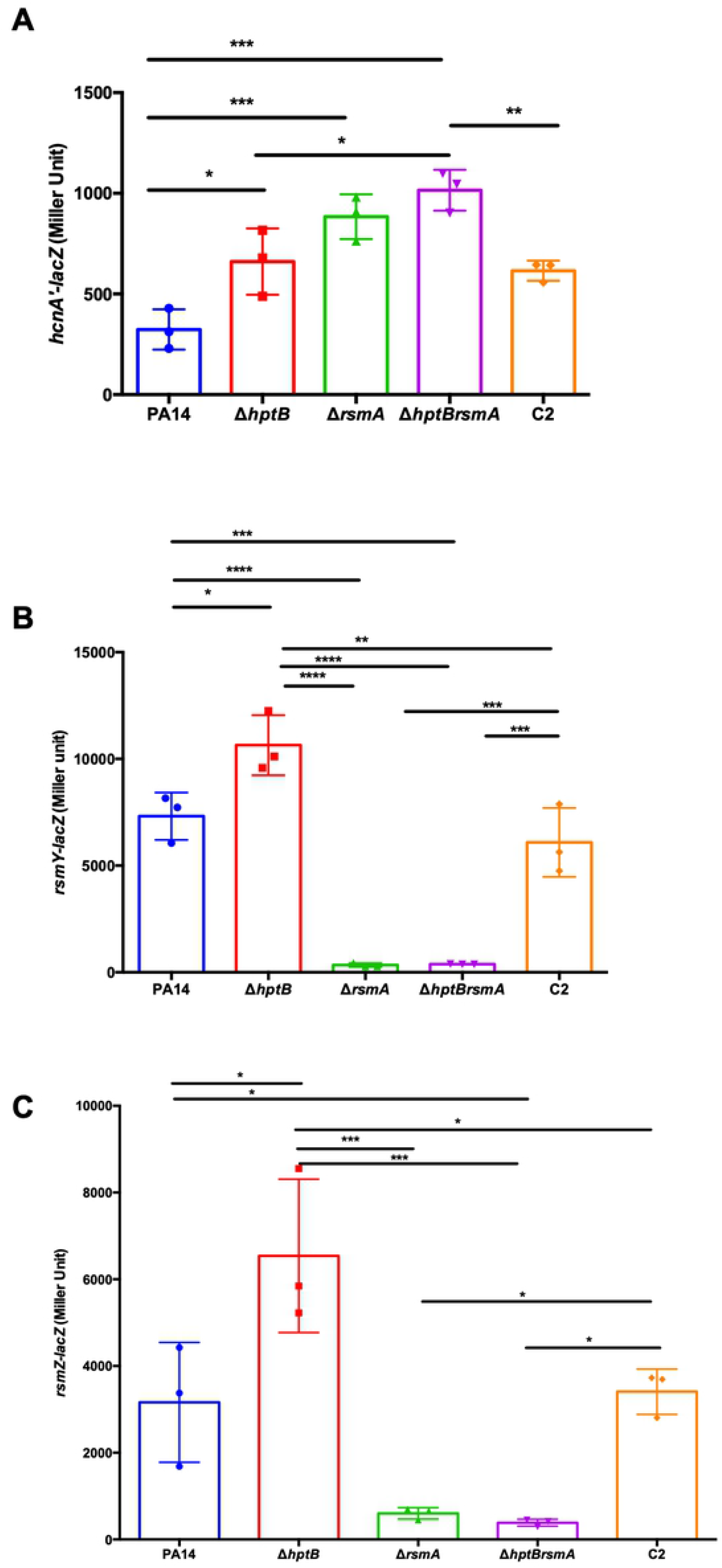
Expression of RmsA-regulated targets in the C2 clone mutant. (A) β-galactosidase activity of *hcnA*-*lacZ*, (B) *rsmY-lacZ* (C) and *rsmZ-lacZ* at 34°C in M9DCAA at 8h for *hcnA-lacZ* and 6h for *rsmY-lacZ* and *rsmZ-lacZ*. One-way ANOVA was performed to compare each strain to each other. (*≤0.05, **≤0.01, ***≤0.001, ****≤0.0001). See **Fig S4** for the growth of each strain.

### The R31S mutation alters RsmA affinity for RsmY

One hypothesis to explain our observations is reduced binding affinity, and thus repression, of RsmA^R31S^ activity by RsmY/Z. We first tested this hypothesis by building a molecular model of the RsmA-RsmY complex. This allowed us to evaluate the atomic-scale impact of the R31S mutation on the structure and binding energy of complex formation between WT RsmA and RsmA^R31S^. Our structural model suggests that the wild-type R31 residue is most likely involved in the stabilization of the U88A89 nucleotide pair located downstream of the conserved GGA motif in RsmY. Replacing the long, charged, and flexible terminal guanidinium arginine moiety for the small and polar hydroxyl group of a serine side chain results in the loss of two key hydrogen bonding interactions between RsmA and RsmY **(Fig. 5)**. In addition, increased local flexibility combined with perturbations in steric contacts and short-/long-range electrostatic interactions result in a total estimated energy penalty contribution of ~18 kcal/mol in variant RsmA^R31S^ relative to wild-type RsmA. This energy penalty most likely results in important binding variability within the protein-ligand ensemble, especially considering that binding free energy (Δ*G*) values of 2-3 kcal/mol are sufficient to impart significant alterations in protein-RNA interactions (27, 28).

**Figure 5.**
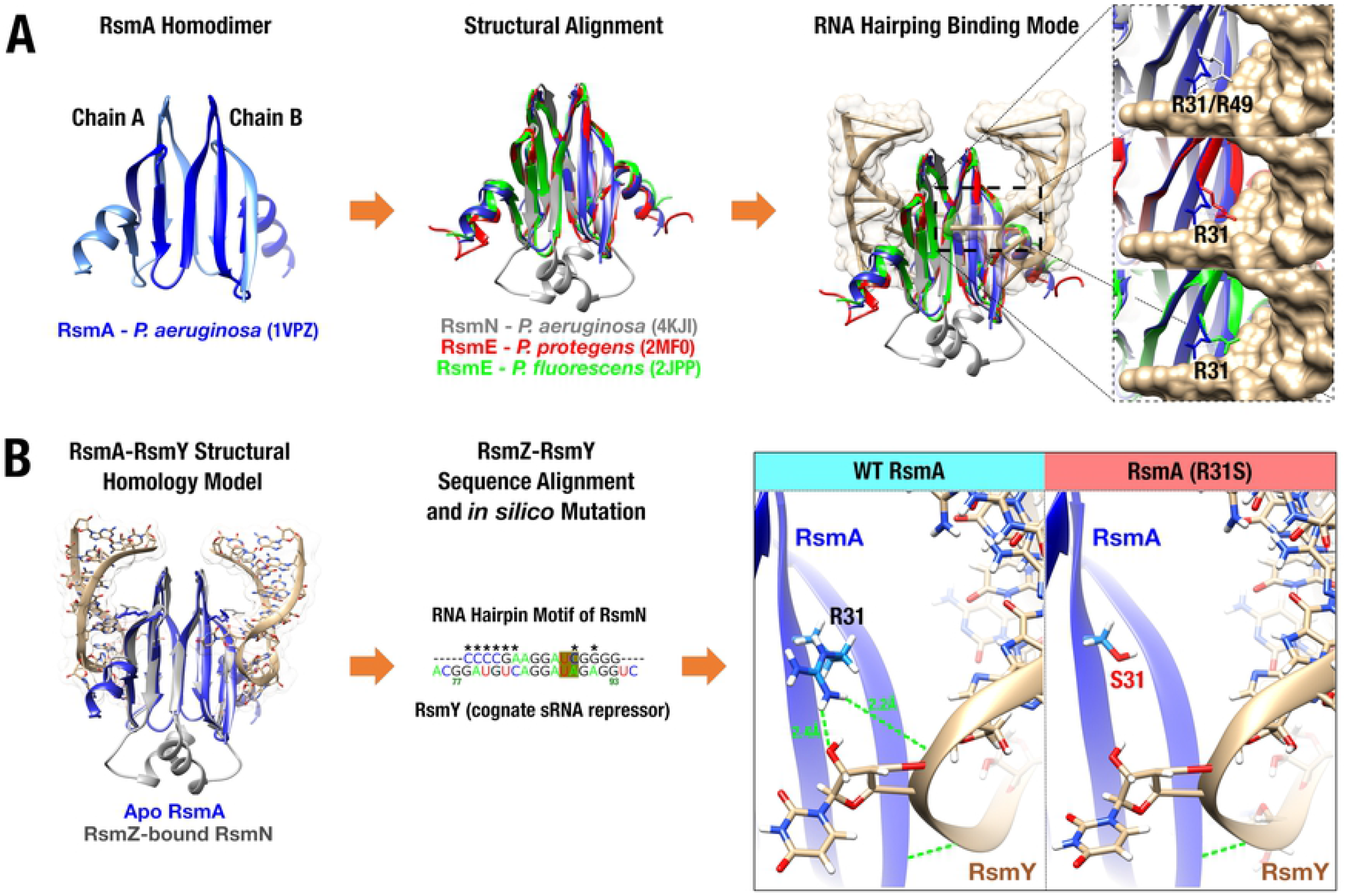
Prediction of RsmA^R31S^ interaction with sRNA RsmY. Structural effects induced by the R31S mutation in post-transcriptional regulator RsmA upon binding to its sRNA repressor RsmY. (A) Structural comparison between RsmA from *P. aeruginosa* (blue, left and middle panel) and three homologous regulators (middle panel): RsmN from *P. aeruginosa* (gray, PDB 4KJI), RsmE from *P. protegens* (red, PDB 2MF0), and RsmE from *P. fluorescens* (green, PDB 2JPP). Overlay illustrates structural conservation of post-transcriptional regulators throughout the *Pseudomonas* genus. Regulator protein homologs also bind RNA targets in a similar fashion, involving the conservation of a structural RNA hairpin motif (gold surface), which triggers repositioning of the conserved R31 side chain (R49 in RsmN) upon RNA binding (right panel). This further emphasizes the role played by the wild-type R31 in stabilization of the U88A89 nucleotide pair located downstream of the conserved GGA motif in RsmY. (B) Structural model of the RsmA-RsmY complex built using the highly homologous RsmN-RsmZ pair as cognate template (gray, PDB 4KJI). *In silico* point mutations (middle panel, nucleotides marked with a star) were introduced to convert the RsmZ hairpin sequence motif into the corresponding putative RsmY hairpin analog bound to RsmA. The right panel shows how the R31S mutation destabilizes the RsmA-RsmY binding complex by abrogating two putative hydrogen bonding interactions (green dashed lines) between the terminal wild-type R31 guanidinium moiety and the U_88_A_89_ nucleotide pair downstream of the conserved GGA motif in cognate RsmY repressor.

We then used electrophoretic mobility shift assays (EMSA) to experimentally challenge this model and investigate interactions between wild-type and RsmA^R31S^ upon RsmY binding **(Fig. 6)**. Our EMSA results show that RsmA binds to RsmY and forms three different protein-RNA complexes, which are reflected by the known binding of several RsmA monomers to multiple binding sites on a RsmY sRNA molecule (29, 30). The RsmY sRNA has seven GGA sites where RsmA can bind. The second, fifth and seventh binding sites are the most determinant for RsmA-RsmY complex formation (29). At tested RsmA concentrations where protein:RNA interactions are detected, both wild-type and RsmA^R31S^ at the concentration of 0.05 μM formed complex 1 **(Fig. 6B)**. However, at higher concentrations, RsmA^R31S^ can only form complex 2. In contrast, the wild-type protein was able to shift RsmY, forming complex 3, which was not observed for RsmA^R31S^ at the same concentration **(Fig. 6A and Fig. S5)**.

**Figure 6.**
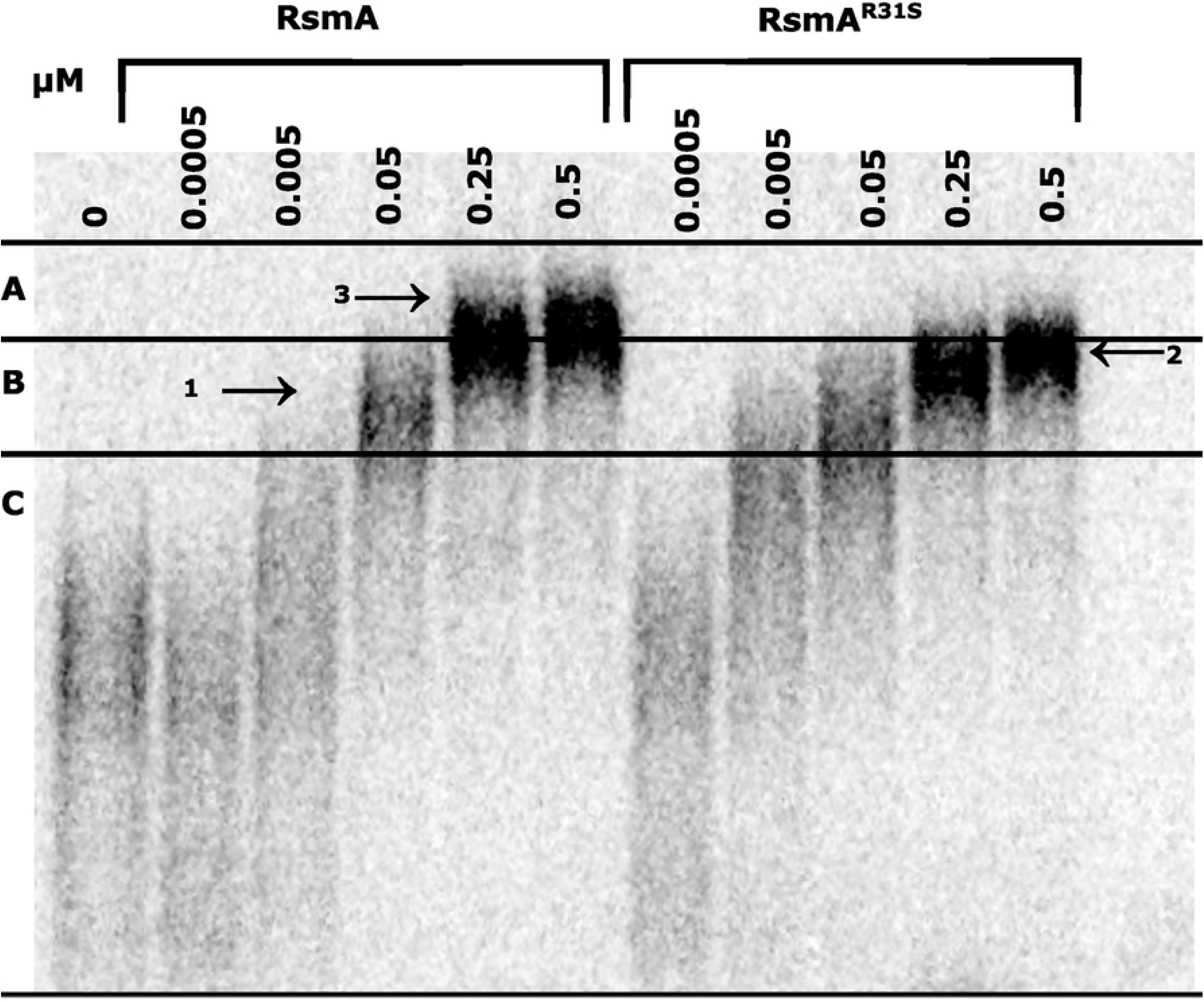
*In vitro* interaction of RsmA and RsmA^R31S^ with RsmY. EMSA of *rsmY* small RNA by wild-type RsmA (WT) and mutated RsmA^R31S^ with 0 to 0.5 μM of each protein. Sections A, B, C represent the delimitation of the boxes used to measure radioactivity for **Fig S6**. Complexes are identified by numbers 1-2-3 and arrows on the EMSA.

To confirm our observations, the radioactive intensity of each lane was measured for each section found in **figure 6 (A-B-C)** to quantify the three complexes. The intensity of each section was then divided by the total intensity of the lane (sum of intensities of sections A, B, C for each concentration) to determine the ratio of RNA for each section **(Fig. S6)**. In section A, complex 3 was found and could be quantified **(Fig. S6A)** while the first and second complexes were quantified in section B **(Fig. S6B)**. Unbound RNAs were quantified in section C **(Fig. S6C)**. These data demonstrate that RsmA^R31S^ does not bind as well as wild-type RsmA to RsmY, which validates our *in silico* model and explains the results we observed for our *in vivo* assays.

## Discussion

Here we used swarming motility as a model social phenotype to better understand the implication of HptB and the Gac/Rsm pathway in surface behavior of *P. aeruginosa*. The post-transcriptional regulator RsmA favors swarming motility and the acute mode of infection while repressing functions involved in the development of chronic infections, such as biofilms. Even though swarming is regulated inversely than biofilm formation, evidence suggests that swarming could play a role in early steps of biofilm development (31, 32). Furthermore, swarming cells are more resistant to antibiotics (33). Our previous report on the swarming-deficient *hptB* mutant demonstrated that flagellar motility and production of a wetting agent are not the only elements essential for swarming motility; using directed evolution of Δ*hptB* under swarming conditions, we found that *lasR*-defective mutants partially regained the ability to swarm (23). Here, we investigated a second group of mutants in the Gac/Rsm pathway that also arise in a Δ*hptB* background during experimental swarming evolution and that are fully rescued in their surface motility. One of these mutants (clone C4) had a modification in the *gacA* gene resulting in a protein with a deficient activity. Indeed, the transcription of both *rsmY* and *rsmZ* in clone C4 are at the same level as in a Δ*hptBgacA.* This finding was not surprising, as we had previously found that loss of *rsmY* and *rsmZ,* which are the primary targets of GacA, completely rescues the swarming defect in a Δ*hptB* background (34). Spontaneous mutations in the *gacA* and *gacS* genes have been previously well-documented in different *Pseudomonas* strains and various growth conditions (35–37), including in the context of swarming evolution experiments (36). The mutation found in *gacA* is located at a well-conserved position (38, 39) in the receiver domain next to the aspartic acid essential for GacS phosphorelay (39). Since the mutation led to a loss-of-function, it could prevent the phosphotransfer by the GacS sensor. Loss of GacA essentially abolishes *rsmY* and *rsmZ* expression, resulting in an increased availability of RsmA and thus increased repression of its multiple mRNA targets leading to an indirect promotion of swarming motility.

The experimental evolution with the Δ*hptB* mutant also selected for a mutation in the *rsmA* gene which led to a restored swarming phenotype. This was completely unexpected since loss of RsmA activity decreases swarming motility **(Fig. 1C)** (24). A spontaneous mutation in that gene has never been reported. The best explanation for this surprising mutation was that RsmA has an altered protein activity caused by this single nucleotide substitution. In contrast with the Δ*hptBrsmA* and Δ*rsmA* mutants, the evolved C2 clone is capable of a wild-type-like swarming, which further supports the observation that the mutation obtained in the evolved clones does not result in a non-functional protein. Those results also concur with *rsmY* and *rsmZ* expression assays: transcription of both sRNAs is lower in Δ*rsmA* and Δ*hptBrsmA* compared to PA14, but clone C2 is similar to PA14. The same observations were also made when looking at the translation of *hcnA*. These results support a model where the emergence of the single nucleotide substitution mutation in *rsmA* results in a modification rather than a loss of protein function.

To better understand how the emergence of a RsmA^R31S^ substitution in the evolved Δ*hptB* strain could rescue its swarming motility phenotype without affecting the expression of *rsmY* and *rsmZ*, we hypothesized that the affinity between RsmA^R31S^ and target RNAs could be impacted. In *Escherichia coli* K12, a R31 substitution of CsrA (RsmA homolog) to alanine subtly affects the *in vivo* regulation of CsrA on target genes or phenotypes (40). However, this effect is not as important as observed with other neighboring residues. R31 is not strictly conserved between bacterial species, although positively charged residues are primarily found at this position (arginine, lysine, histidine) (40, 41) **(Fig. S7)**. Residue R44 is involved in RNA interaction and R31 most likely plays an important accessory role impacting binding affinity and/or ligand discrimination since it appears to stabilize the U_88_A_89_ nucleotide pair located downstream of the conserved GGA motif in RsmY **(Fig. 5)**. This positively charged residue is solvent-exposed and its interaction with RNA is mediated by two hydrogen bonds involving its terminal guanidinium moiety (40, 41). In contrast, the mutation in C2 introduces a serine instead of an arginine, leading to a small and polar residue unable to maintain these interactions, therefore affecting RNA binding affinity and/or discrimination.

We confirmed that RsmY affinity for RsmA with the R31S substitution is modified due to different RsmY mobility shift when interacting with either RsmA or RsmA^R31S^ in EMSA experiments **(Fig. 6)**. The most probable interpretation is that the loss of interaction between the R31 residue and RNA reduces affinity for some binding sites, notably with GGAUA, such as in RsmY, as opposed to other sites. The complexes between sRNA and RsmA are the result of the multiple RsmA molecule binding to the different GGA motifs of RsmY (29). Even though both RsmA and RsmA^R31S^ are capable of interacting with RsmY and forming Complex 1, only the wild-type protein can form Complex 3 which most likely represents a higher capacity to bind RsmY molecules compared to RsmA^R31S^. Indeed, this was confirmed when we looked at the radioactivity signal of the protein-RNA complexes; clearly, RsmA^R31S^ does not bind as well as WT RsmA to RsmY **(Fig. S6A)**. Affinity between RsmA^R31S^ and RsmY shows that the inhibition by RsmY is not as efficient, which supports the fact that the C2 clone exhibits similar activity as the wild-type protein **(Fig. 4B),** but different than Δ*rsmA* and Δ*hptBrsmA.*

Our results indicate that RsmA inhibits, directly or indirectly, an unknown repressor impacting swarming motility. However, this repressing factor is still unknown, and it is thus not yet possible to test its activity with RsmA and RsmA^R31S^ *in vitro*. RsmY has many RsmA binding sites and displays secondary structures with multiple stem-loops with RsmA binding sites (29). Depending on the different mRNAs that are controlled by RsmA, the number of available binding sites and/or secondary structures could probably affect the binding capacity between the protein and target RNAs. The RsmA:mRNA interaction of these RsmA-controlled mRNAs implicated in swarming motility could be less impacted by the mutation R31S than RsmY, due to their sequence and structure, and then explain the rescue of swarming in the Δ*hptB* background. Also, additional elements such as chaperone Hfq, which can bind some mRNA transcripts that associate with RsmA, could contribute to RNA binding and were missing in our *in vitro* experiments (42). However, our data strongly support the fact that a modified RsmA binds RsmY less efficiently, impacting its inhibitory effect, which further explains the swarming rescue of the C2 clone.

We wanted to understand why the swarming population of an evolved Δ*hptB* mutant selected for a spontaneous mutation in RsmA. HptB can inhibit *rsmZ* expression under swarming conditions independently of GacA (13) and also inhibits *rsmY* and *rsmZ* by indirectly influencing GacA. We can then hypothesize that a Δ*hptB* favours the inhibition of RsmA by increasing expression of *rsmZ* and *rsmY* which could explain its lack of swarming. The RsmA mutation in the C2 evolved clone most likely results in lower affinity for RsmY and RsmZ, compensating for the Δ*hptB* mutation. The mutated RsmA version has lower *rsmZ* and *rsmY* expression than Δ*hptB* mutants. This could explain the selection of a spontaneous mutant in our directed evolution experiment and thus the benefit of having a mutation that modifies RsmA activity. Additionally, although arginine is primarily found at position 31, we found two naturally occurring RsmA variants at this position in *P. aeruginosa* strains using BLAST (NCBI) (R31G and R31C). We aligned these sequences (and other sequences of RsmA homologs in different strains) to PA14 RsmA sequence in a multiple sequence alignment to observe the variation found at position 31 (**Fig. S7).** This can indicate that naturally occurring RsmA can tolerate R31 replacements, although R31 is largely favored for optimal activity. Understanding this mutation could help identify the unknown element(s) necessary for swarming motility.

Previous experimental evolution on wild-type PA14 strain under swarming conditions did not reveal a mutation in *rsmA* (43, 44). Also, passaging Δ*hptB* in liquid cultures did not result in a rescue of swarming motility (23). It is likely that the use of swarming conditions and a *hptB* mutant background have selected for the mutation in *rsmA*. Our results show the central role of RsmA in swarming regulation. Decreasing RsmA repression by its cognate sRNAs was the only evolved way to relieve sRNA-mediated repression of swarming in Δ*hptB* mutant. We could achieve it through selective pressure forcing the Δ*hptB* mutant to swarm; something that would have not been obtained if we had adopted a classical transposon mutagenesis screening approach where only loss-of-function mutants can result. We confirmed the role of RsmA as a master switch between bacterial acute (motile lifestyle) and chronic infection (sessile lifestyle) and identified a residue that is critical for its activity. Knowing this, we could now modulate its function by permitting a better adaptation to its environment. Even though RsmA regulates secondary metabolites, it could act as a target to inhibit biofilm formation.

## Materials and methods

### Bacterial Strains and culture conditions

*Pseudomonas aeruginosa* strain PA14 was used in this study (45). Details on the strains are found in **Table S1**. Conditions used for the directed evolution under swarming conditions was previously described (23). The bacteria were grown in Tryptic Soy Broth (TSB) (Difco) at 37°C in a TC-7 roller drum (New Brunswick) at 110 rpm unless otherwise specified. Swarming assays have been performed as previously described (46). Swarming plates containing 0.5% agar were prepared and 5 μl of a bacterial suspension at OD_600_=3.00 were inoculated. The plates were incubated at 34°C in bags for 20h or as indicated. For the swarming complementation experiment, the swarming plates were supplemented with 125 μg/mL of tetracycline. Pictures were taken using a Lumix DMC-ZS60 camera (Panasonic). OD_600_ measurements were taken using a Nanodrop ND-1000 (Thermo Fisher Scientific).

### Sequencing of the whole genome DNA

The whole genome sequencing of C2 and C4 was obtained as previously described (23). The C2 and C4 clones generated 2,862,314 and 3,254,964 reads respectively covering a 6M bp genome. Mutations were confirmed by PCR amplification (**Table S2**). Purified PCR fragments were sent to Institut de Recherches Cliniques de Montréal (Montréal, Canada) for Sanger Sequencing. RsmA/CsrA variants from other *P. aeruginosa* strains and other bacterial species were obtained by BLAST (NCBI https://www.ncbi.nlm.nih.gov/) and sequences aligned was performed through Clustal Omega 1.2.3 on Geneious Prime 2020.0 (Biomatters).

### Construction of markerless *rsmA* and *gacA* mutants

The *rsmA* gene (PA14_52570) was deleted at more than 95% using a two-step allelic exchange method (47). All primers are described in **Table S2**. The upstream region was amplified using RsmA-L-F-EcoRI and RsmA-L-R-homRsmA primers. The downstream fragment was amplified using primers RsmA-F-F and RsmA-R-R-HindIII. And the overlapping PCR was amplified using RsmA-L-F-EcoRI and RsmA-R-R-HindIII. The obtained fragment was ligated into pEX18-Ap. The obtained vector (pSR09) was transformed in conjugative *E. coli* SM10 and selected on Lysogeny Broth (LB) Miller’s agar (Alpha Biosciences) containing 100 μg/ml of carbenicillin. The allelic exchange was performed in PA14 and Δ*hptB* by conjugation. Clones were selected on carbenicillin 300 *μ*g/ml and triclosan 25 *μ*g/ml. The double recombination was performed on no salt LB agar with the addition of 10% sucrose.

The same method was used for the deletion of the *gacA* (PA14_30650) gene. Primers are listed in **Table S2**. The upstream region has been amplified using primers FJP_UP_gacA_pEX_For and FJP_UP_gacA_pEX_Rev. The downstream fragment has been amplified using FJP_DN_gacA_pEX_For and FJP_DN_gacA_pEX_Rev primers. The overlapping PCR was amplified with FJP_UP_gacA_pEX_For and FJP_DN_gacA_pEX_Rev. The obtained fragment was inserted in pEX18-Ap. The plasmid obtained (pFJP19) was transformed in SM10 and selected on LB agar with 50 μg/ml carbenicillin. Allelic exchange in PA14 and Δ*hptB was* performed as described (47).

### Construction of strains with gene expression reporters

Two-partnered conjugation with SM10 containing *pCTX-rsmY* or pCTX-*rsmZ* was performed with *P. aeruginosa* strains. Clones were selected on 125 *μ*g/ml tetracycline and 25 *μ*g/ml triclosan. Plasmid pME3826 was transformed by electroporation as described previously (48). Clones were selected on LB agar containing 125 μg/ml of tetracycline.

### B-galactosidase assays

For the expression of *rsmY-lacZ, rsmZ-lacZ,* and *hcnA’-lacZ,* strains were inoculated in TSB from frozen stock and incubated as previously described with tetracycline when needed. Overnight cultures were diluted in fresh TSB or M9DCAA modified medium (46) as specified at OD_600_=0.5. The cultures were incubated at 34°C or 37°C in a TC-7 roller drum (New Brunswick) at 110 RPM. β-galactosidase activity was measured as previously described (49). Measurements at 420 nm were performed using a Cytation 3 Multiplate Reader (Biotek). Experiments were performed using three biological replicates and were repeated at least twice. Prism 6 (GraphPad) was used for statistics.

### Complementation experiments

The *rsmA* gene was amplified from gDNA of PA14 WT and the evolved C2 clone using primers For_Comp_rsmA_EcoRI_FJP and Rev_Comp_rsmA_HindIII **(Table S2)**. The amplified fragment was inserted into pUCP20 using HindIII and EcoRI restriction sites. The obtained clones were isolated on LB agar with 100 *μ*g/ml carbenicillin. The obtained plasmid was electroporated into PA14Δ*rsmA rsmY-lacZ* and clones were selected with 250 *μ*g/ml carbenicillin. PA14 *rsmY-lacZ* with pUCP20 was used as a control.

Complementation of Δ*hptB* and *rsmA*::MrT7 with wild-type *rsmA* was achieved by amplifying the sequence of *rsmA* from genomic DNA of PA14 using the For_Comp_rsmA_EcoRI_FJP and Rev_Comp_rsmA_HindIII primers. The obtained fragment was inserted in the pUCP26 vector by digestion using EcoRI and HindIII Fast Digest restriction enzymes (Thermo Fisher Scientific). The plasmid obtained (pFJP18) was transformed in DH5α and selected on LB agar with tetracycline 15 *μ*g/ml. The construct was verified by digestion with EcoRI and HindIII restriction enzymes (Thermo Fisher Scientific) and PCR. The pFJP18 plasmid was then electroporated into PA14Δ*hptB* and *rsmA*::MrT7 along with empty pUCP26 vector and clones were selected with tetracycline 125 *μ*g/ml. Four independent clones were tested for their swarming phenotypes.

### Purification of RsmA

For purification of RsmA-WT-6xHis and RsmA^R31S^-6xHis, plasmids pET29a(+)-RsmAH6 or pET29a(+)RsmA^R31S^ were used respectively to produce the protein. Plasmid pET29a-RsmA^R31S^-H6 was created by inserting a synthesized mutated version of RsmA^R31S^ without start and stop codon into NdeI and XhoI restriction sites (BioBasic) of pET29a(+). Plasmids were transformed in BL21 (DE3).

Purification was performed as previously described with slight modifications (34). The BL21(DE3) strain containing either pET29a(+)-RsmA-H6 or pET29a(+)-RsmA^R31S^-H6 was grown overnight in TSB containing kanamycin 50 *μ*g/ml. Overnight cultures were diluted 1:1000 and grown to exponential phase in LB containing kanamycin 50 μg/ml at 37°C with shaking at 250 rpm. Cells were induced by adding IPTG at a final concentration of 1 mM and grown for an additional 4h. The culture pellet was suspended in a buffer containing 0.5 M NaCl, 20 mM NaH_2_PO_4_, 20 mM Tris-HCl pH7.5 and 2% imidazole. To purify the protein, a HisTrap HP 5 ml column (GE Healthcare) was used. Purification was performed using a solution of 0.5 M NaCl, 20 mM NaH_2_PO_4_, 20 mM Tris-HCl pH 7.5. To elute the His-Tagged protein, a solution of 0.5 M NaCl, 20 mM NaH2PO4, 20 mM Tris-HCl pH 7.5 and 0.5 M imidazole was applied using a 2-50% gradient in 100 minutes, completing with up to 100% imidazole solution in 15 minutes using an ÄKTA FPLC system (GE Healthcare). Fractions containing the protein were pooled and concentrated using an Amicon with a 3 KDa cut-off (Millipore). The protein was conserved in 10 mM Tris-HCl pH7.63 and 33% glycerol. A Bradford assay (Bio-Rad) was performed to determine protein concentration. Confirmation was performed using a Western-Blot with anti-His antibodies (Mouse antibodies to 6His-peptide) (Meridian Life Science) and Coomassie blue coloration to confirm protein purity.

### Electrophoretic mobility shift assays of *rsmY* and RsmA/RsmA R31S (C2)

The *rsmY* small RNA template with a T7 RNA polymerase promoter sequence was produced as previously described (34). The fragment was amplified using 5’-rsmY and 3’-rsmY primers (**Table S2**) and a nested PCR was then performed using *3’-rsmY* nested as the reverse primer. Purification was done using a FavorPrep^TM^ Gel/ PCR purification kit (Favorgen). The negative control was the Ykok riboswitch of *Halorhodospira halophila* SL1 (NC_008789.1). The sequence is coded at the position 1425650-1425831 on the positive strand of the genome.

For *in vitro* radioactive transcription, the templates were added to the transcription reaction containing 0.5 μl of Ribolock RNAse inhibitor at 40U/μl (Thermo Fisher Scientific), 1 μl of pyrophosphatase at 5 *μ*g/ml (Roche), 4 *μ*l of 1 mg/mL T7 RNA polymerase and 20 *μ*l of transcription buffer 5X (Hepes pH7.5 400 mM, MgCl_2_ 120 nM, DTT 200 mM and spermidine 10 mM). For nucleotides, 5 *μ*l of 100 mM of ATP, GTP and CTP were added. For UTP, 1 *μ*l of 2 mM of non-radiolabelled and 5 μCi [α^32^P]-UTP was added to the mix. The final volume was 100 μl per reaction. The reaction was 3h at 37°C. A DNase I RNAse free 2000 U/ml (NEB) treatment was done using 1 μl for 15 minutes at 37°C. The resulting RNA was purified on an 8% 19:1 acrylamide:bisacrylamide 8M urea denaturing PAGE and resuspended in 250 μl of RNAse-free water.

For the EMSA assay, various concentrations of each protein were mixed to 2 μl of radio-labelled RNA in a reaction described in Jean-Pierre *et al.* (34) containing 20 mg of non-specific t-RNA competitor, 10 mM Tris-HCl pH7.5, 10 mM MgCl_2_,, 50 mM NaCl, 50 mM KCl, 5 mM DTT. Negative control was done in the same condition containing no protein but containing glycerol and Tris-HCl 10 mM pH7.5 (the protein dilution buffer). The reaction was incubated 30 minutes at 37°C. The mixture was added mixed to 6X loading buffer containing (40% sucrose, 0.05% xylene cyanol, 0.05% bromophenol blue) and loaded on an 8% (29:1) native polyacrylamide native gel using Tris-Borate EDTA (TBE) as the running buffer. The gel was run at room temperature for 6h at 150V. The gel was scanned using a Typhoon PhosphorImager FLA9500 (GE Healthcare) and ImageQuant software was used for analysis of the image. The gel was repeated twice with similar result.

### Molecular modeling of the WT and R31S RsmA-RsmY complexes

To investigate atomic-scale contributions of the R31S mutation in RsmA, structural alignments were first performed between apo and holo forms of RsmA, RsmE, and RsmN homologs using PDB entries 1VPZ, 4KJI, 2MF0, and 2JPP in UCSF Chimera 1.14 (50). Since regulators bind their respective RNA ligands through a conserved hairpin RNA motif, the RsmY ligand was modeled from the experimental structure of the RsmZ analog bound to RsmN (PDB 4KJI). RNA sequence alignment was performed using the Needleman-Wunsch algorithm in package BioLabDonkey 1.9-17. We first identified RsmY/Z hairpin motif sequence identity, followed by mutational transposition of RsmY nucleotides in the RsmZ structural template. Once the molecular sRNA hairpin motif was created, the RsmA^R31S^ mutant was built in a similar fashion by replacing Arg31 with Ser31. A physiological pH value of 7.4 was applied to assign protonation states of charged amino acids, with pKa values predicted according to parameters reported by (51). Rotameric positions and RsmA-RsmY refinement for WT and R31S complexes was optimized by performing molecular dynamics simulations (500 ps, 298K) under explicit solvent conditions with a water density of 0.997 g/ml and pressure density stabilized by the Manometer1D tool. This protocol enables a rescaling factor to be applied over the entire MD cell (cuboid shape) to maintain constant pressure during simulations. The unit cell was extended 10 Å with solute on each side of the system, and ion concentration was set as a mass fraction of 0.9% NaCl to emulate physiological conditions. The simulation time step ran at 2×1.25 fs in periodic boundary conditions using particle-mesh Ewald (PME) and 8.0 Å cutoff for non-bonded real space forces. The CorrectDrift algorithm was applied to prevent solute molecules from diffusing around and crossing periodic boundaries. A final energy minimization step was performed after refinement of the RNA-protein complexes. Building of the RsmY RNA hairpin motif, structural refinement, molecular dynamics simulations (MD), and energy minimization steps for all molecular systems were performed using YASARA-Structure 19.12.14 (52). The YASARA2 force field was applied for refinement, solvation, and MD simulations of RsmA-RsmY complexes (53). RNA-protein interface analysis of all complexes was also performed using the MolDock scoring function provided by the Molegro Virtual Docker suite, version 6.0 (54).

### Stability and energy contributions of the R31S mutation in RsmA

The structural and energetic effects caused by the R31S mutation were assessed by evaluating unfavorable torsion angles and investigating the local structural environment of the mutated position using CUPSAT (55). We also used DUET to perform mutational analysis based on energy function calculations (56). Analysis and prediction of protein stability changes upon mutation was also performed by Normal Mode Analysis using DynaMut (57). Finally, Mupro was used to calculate neural networks that compute the effects of mutation from sequence and structure predictions (58).

## Acknowledgments

We thank Charles Morin for construction for the Δ*gacA* and Δ*hptBgacA* mutants and for helpful discussions and Sabrine Najeh for providing labelled YkoK riboswitch control. We also gratefully acknowledge the technical support provided by Myriam Létourneau and Émilie Boutet for the RsmA purification and EMSA assays, respectively. SR obtained PhD scholarship from the Fonds de recherche du Québec – Nature et technologies (FRQNT) and also from the Fondation Armand-Frappier de l’INRS.

## Supplemental Figures

**Figure S1. Growth of the C4 clones and related strains.** The growth of (A) evolved clone C4 containing a *rsmY-lacZ* reporter and associated strains, (B) evolved clone C4 and associated strain containing *rsmZ-lacZ*, in TSB at 37° C at 5h of incubation. One-way ANOVA was used to compare each strain to each other. Prism 6 (GraphPad) was used for statistics (*≤0.05, **≤0.01, ***≤0.001, ****≤0.0001).

**Fig S2. Swarming of Δ*hptB* mutant when complemented with a plasmid-borne *rsmA*.** Swarming motility of Δ*hptB* and *rsmA^-^* with pUCP26 or pFJP18 (pUCP26-RsmA) on M9CAA with added tetracycline.

**Figure S3. Growth of the complementation strains.** Growth of the strains for the complementation of RsmA in TSB at 37°C at 6h. One-way ANOVA was used to compare each strain with each other strains (*≤0.05, **≤0.01, ***≤0.001, ****≤0.0001).

**Figure S4. Growth of the C2 clones and related strains.** Growth of the strains with the (A) *hcnA*-lacZ reporter (B) *rsmY-*lacZ and (C) *rsmZ-*lacZ in M9DCAA medium at 34°C 8h for *hcnA-lacZ* and 6h for *rsmZ-lacZ* and *rsmY-lacZ*. One-way ANOVA was used to compare each strain with each other strains (*≤0.05, **≤0.01, ***≤0.001, ****≤0.0001).

**Figure S5. Binding of RsmA and RsmA^R31S^ to negative control.** EMSA YkoK riboswitch RNA by wild-type RsmA (WT) and mutated RsmA^R31S^ with 0 to 0.5 μM of each protein. The YkoK riboswitch from *Halorhodospira halophila* SL1 was used as a negative control. This RNA is not known to be regulated by RsmA. Although, the negative control shifts at high protein concentrations, the RsmY shift was achieved at 100 times lower protein concentrations than YkoK riboswitch. There is a GGA in the sequence of the riboswitch that could explain the slight shift.

**Figure S6. Quantification of the binding between RsmY and RsmA^R31S^.** Graphic representation of the binding of RsmA and RsmA^R31S^ to RsmY(A) when forming Complex 3, (B) when forming Complex 1 and 2, and (C) when unbound. The ratio of intensity of each section is represented on total intensity for each concentration. See **Fig. 6** for each complex and section where radioactivity is measured **(Fig. 6A-B-C)**. The intensities shown are the mean of the results from two independent gels having similar results and error bars are the standard deviation

**Figure S7. Comparison of CsrA sequences from *P. aeruginosa* strains and other species.** CsrA protein sequences were obtained through BLAST (NCBI) of the RsmA sequence from PA14 (higlight in yellow), and by modify the R31 residue. CsrA from *P. aeruginosa* with 100% identity and modified versions at R31 residue were aligned with other species CsrA with a modified R31 residue using Clustal Omega 1.2.3.

## Supplementary Tables

**S1 Table. Strains used in this study**

**S2 Table. Primers used in this study**

